# Degeneration and Impaired Resilience of Skull Bone and Hematopoietic Bone Marrow

**DOI:** 10.1101/2025.10.02.679940

**Authors:** Yang Yang, Yonggang Fan, Sanyam Jain, Zhangfan Ding, Hanyu Liu, Allison L Horenberg, Haiyang Sun, Mara Sophie Killer, Abhishek Chandra, Rinkoo Dalan, Manju Chandran, Lynn Yap, Junyu Chen, Navin Kumar Verma, Christine Cheung, Makrand V Risbud, Aline Bozec, Kristijan Ramadan, Martine Cohen-Solal, Warren L Grayson, Saravana K Ramasamy, Anjali P Kusumbe

**Affiliations:** Tissue and Tumor Microenvironments Lab, Cancer Discovery and Regenerative Medicine Program, Lee Kong Chian School of Medicine, Nanyang Technological University, Singapore, Singapore; Multidisciplinary Institute of Ageing (MIA-Portugal), University of Coimbra, Coimbra, Portugal; Lee Kong Chian School of Medicine, Nanyang Technological University, Singapore, Singapore; Department of Biomedical Engineering, Johns Hopkins University School of Medicine, Baltimore, MD, USA; Translational Therapeutics & Regenerative Engineering Center, Johns Hopkins University School of Medicine, Baltimore, MD, USA; Department of Materials Science and Engineering, Johns Hopkins University, Baltimore, MD, USA; Department of Chemical & Biomolecular Engineering, Johns Hopkins University, Baltimore, MD, USA; Institute for NanoBioTechnology, Johns Hopkins University, Baltimore, MD, USA; Department of Rheumatology and Reference Center for Rare Bone Diseases, Hospital Lariboisière, Paris, France; Inserm U1132 Bioscar, Université Paris Cité, Centre Viggo Petersen Hôpital Lariboisière, APHP.Nord, 75010 Paris, France; The MRC Weatherall Institute of Molecular Medicine, John Radcliffe Hospital, University of Oxford, Oxford, OX3 9DS, UK; Cancer Discovery and Regenerative Medicine Program, Lee Kong Chian School of Medicine, Nanyang Technological University, Singapore; Department of Internal Medicine 3 — Friedrich-Alexander-University Erlangen-Nürnberg (FAU) and Universitätsklinikum Erlangen, Erlangen, Germany; Deutsches Zentrum Immuntherapie (DZI), Friedrich-Alexander-University Erlangen-Nürnberg (FAU) and Universitätsklinikum Erlangen, Erlangen, Germany; Department of Orthopaedic Surgery, Sidney Kimmel Medical College, Thomas Jefferson University, Philadelphia, PA, USA; Graduate Program in Cell Biology and Regenerative Medicine, Thomas Jefferson University, Philadelphia, PA, USA; Institute of Molecular and Cell Biology, Agency for Science, Technology and Research, Singapore; State Key Laboratory of Oral Diseases, National Center for Stomatology, National Clinical Research Center for Oral Diseases, West China Hospital of Stomatology, Sichuan University, Chengdu 610041, China; National Heart Research Institute Singapore, National Heart Centre Singapore, Singapore, 169609, Singapore; Osteoporosis and Bone Metabolism Unit, Department of Endocrinology, Singapore General Hospital, Singapore, Singapore; Duke-NUS Medical School, Singapore, Singapore; Department of Endocrinology, Tan Tock Seng Hospital, Singapore, Singapore; Department of Physiology and Biomedical Engineering, Mayo Clinic, Rochester, MN 55905, USA; Robert and Arlene Kogod Center on Aging, Mayo Clinic, Rochester, MN 55905, USA; Department of Biochemistry and Molecular Biology, Mayo Clinic, Rochester, MN 55905, USA; State Key Laboratory of Oral Diseases, National Center for Stomatology, National Clinical Research Center for Oral Diseases, Department of Head and Neck Oncology, West China Hospital of Stomatology, Sichuan University, Chengdu, China

**Keywords:** skull, bone marrow, ageing, blood vessels, haematopoietic

## Abstract

Bone marrow health is central to transplantations, blood formation, and cancer progression. However, the bone marrow niche deteriorates with age, impairing haematopoietic stem cell function. Contrary to a recent report^1^ suggesting skull marrow resists ageing, our multi-laboratory investigation reveals the opposite: the skull marrow is among the vulnerable sites of age-related decline. Ageing skull niches consistently show loss of mesenchymal and osteoprogenitors, suppression of angiogenic and lymphatic programs, adipocyte accumulation, vascular senescence, DNA replication stress, mitochondrial dysfunction, cellular senescence, and heightened inflammation. Proteomic profiling further highlights this vulnerability, demonstrating that vertebral niches—unlike the skull—are relatively spared from these ageing hallmarks. Together, these convergent datasets overturn the notion of skull-specific resilience and instead establish the skull marrow as a fragile, degenerating environment. These findings redefine marrow ageing and highlight the skull as a critical, clinically relevant target for sustaining blood and immune health and reducing vulnerability to haematological disease.

## Introduction

The bone marrow niche progressively deteriorates with age, leading to profound impairments in haematopoietic stem cell (HSC) and skeletal stem cell function^2–4^. Bone marrow niche is composed of highly multicellular complex cell types that interact to maintain HSC function and bone haemostasis^5–10^. Age-associated changes include decline of mesenchymal support, bone loss, depletion of osteoprogenitors, vascular remodelling, increased adipogenesis, immune modulations, heightened inflammatory signalling, accumulation of DNA damage, cancer progression and skewing of HSCs toward the myeloid lineage^11–15^. These changes underpin the decline in regenerative capacity and haematopoietic competence observed in ageing.

Recent work has suggested that skull bone marrow functions as an “expanding and resilient” haematopoietic reservoir, uniquely spared from ageing^1^. If substantiated, such a finding would represent a major shift in current concepts of haematopoietic ageing. In our independent, multi-laboratory analyses employing high-resolution 3D and light-sheet imaging, together with unbiased single-cell transcriptomics and proteomics, we did not find evidence supporting this proposed resilience of skull bone marrow.

We find that the skull bone and bone marrow niche is not spared but undergoes profound age-related degeneration. With ageing, the skull exhibits marked depletion of mesenchymal and skeletal progenitors, loss of angiogenic and lymphatic programs, increased adipocyte accumulation, and elevated levels of proteins associated with DNA damage responses, replication stress, cellular senescence, and inflammatory signalling. Importantly, when directly compared with vertebrae—the principal haematopoietic reservoir and the clinically relevant source for bone marrow transplantation^16,17^—the skull displays accelerated impairment, rather than resilience. These findings are consistent with the broader literature documenting skeletal-site–specific vulnerability of the marrow niche and its critical dependence on bone marrow niche integrity for haematopoietic maintenance.

We also note several methodological concerns in their report^1^ that limit the reliability of its conclusions. Imaging was performed on uncleared whole-mount skulls using confocal microscopy, which is inherently restricted in imaging depth and not well suited for resolving marrow architecture and composition. As a result, the data presented as “skull bone marrow” do not capture the bona fide bone marrow structures. Endomucin is used throughout their study^1^ as a vascular marker to assess age-dependent changes. However, their images show aberrant nuclear localization, which contradicts its established cell-surface expression. Notably, previous *in vivo* labelling of skull vessels by the same lab did not show nuclear accumulation^18^, suggesting that the current staining is artefactual and calls into question the reliability of their vascular imaging data.

Moreover, immunofluorescence images for Endomucin, a transmembrane glycoprotein and endothelial marker^19,20^, show nuclear staining patterns, raising further concerns about the accuracy of the reported vascular architecture^1^. Beyond imaging, the publicly available GEO datasets are not clearly linked to experimental groups described in the manuscript, display inconsistencies in labelling, and include discrepancies in the protocols for HSPC isolation compared with those reported in the paper^1^. Collectively, these methodological limitations substantially weaken the robustness of the study and call into question the claim that skull marrow constitutes a uniquely resilient haematopoietic reservoir^1^.

Our multi-laboratory investigation provides consistent evidence that skull bone marrow represents a vulnerable and degenerating niche during ageing. The notion of skull-specific protection is therefore not supported by our findings and warrant reconsideration. Ageing skull niches reproducibly display hallmarks of ageing. Proteomic profiling further underscores this vulnerability, showing that vertebral niches—unlike the skull—are comparatively protected from several ageing hallmarks. Moving forward, studies of marrow ageing will benefit from rigorous, cross-skeletal comparisons, unbiased cellular and molecular analyses, and transparent methodological practices. Such comprehensive approaches are essential to accurately define the spatial heterogeneity, vulnerability, and functional decline of haematopoietic niches across the skeleton.

## Results

### Age-Associated Impairment of the Skull Bone Marrow Niche and HSC Aging

To examine age-associated remodelling of the skull bone marrow niche, we reanalysed the publicly available single-cell RNA-seq datasets from their study^1^ (GSE275179) (Fig. 1a). Our unbiased comparison of young (YS) and old skull (OS) bone marrow revealed a pronounced decline in niche-supportive compartments in old skulls, including reductions in mesenchymal stem/progenitors (MSPCs), osteoprogenitors, pericytes and endothelial cells (ECs) (Fig. 1b, c). Analyses of specific subsets of *Vcam1⁺* sinusoidal ECs also revealed decline in both old skull and old femurs (Extended Data Fig. 1a). GO term enrichment indicated that these cellular losses were accompanied by transcriptional alterations characteristic of canonical ageing hallmarks, including cellular senescence, impaired telomere maintenance, and reduced stem cell proliferation (Fig. 1d). The aged skull niche further showed downregulation of angiogenic (*Emcn*, *Cdh5*, *Angpt2*, *Egfl7*, *Tek*) and lymphangiogenic (*Prox1*, *Flt4*, *Vegfc*) and reduced expression of endothelial-derived HSC maintenance factors, alongside upregulation of senescence (*Cdkn1a*, *Cdkn2a*, *Trp53*) and inflammatory pathways (*Il6*, *Tnf*) (Fig. 1e, f; Extended Data Fig. 1b). HSPCs from old skulls mirrored these changes with Gene set enrichment analysis of HSPCs demonstrating activation of senescence programs, DNA damage responses, and telomere erosion (Fig. 1g). To directly assess cellular senescence, we used p16 td-Tomato reporter mice, a well-established *in vivo* model for detecting senescent cells. This revealed the presence of senescent ECs in the skulls of old mice (Fig. 1h). Collectively, these findings indicate that the aged skull bone marrow undergoes profound structural and transcriptional decline, displaying hallmarks of accelerated niche and stem cell ageing—contrasting with the suggested model of resilience.

**Fig. 1:**
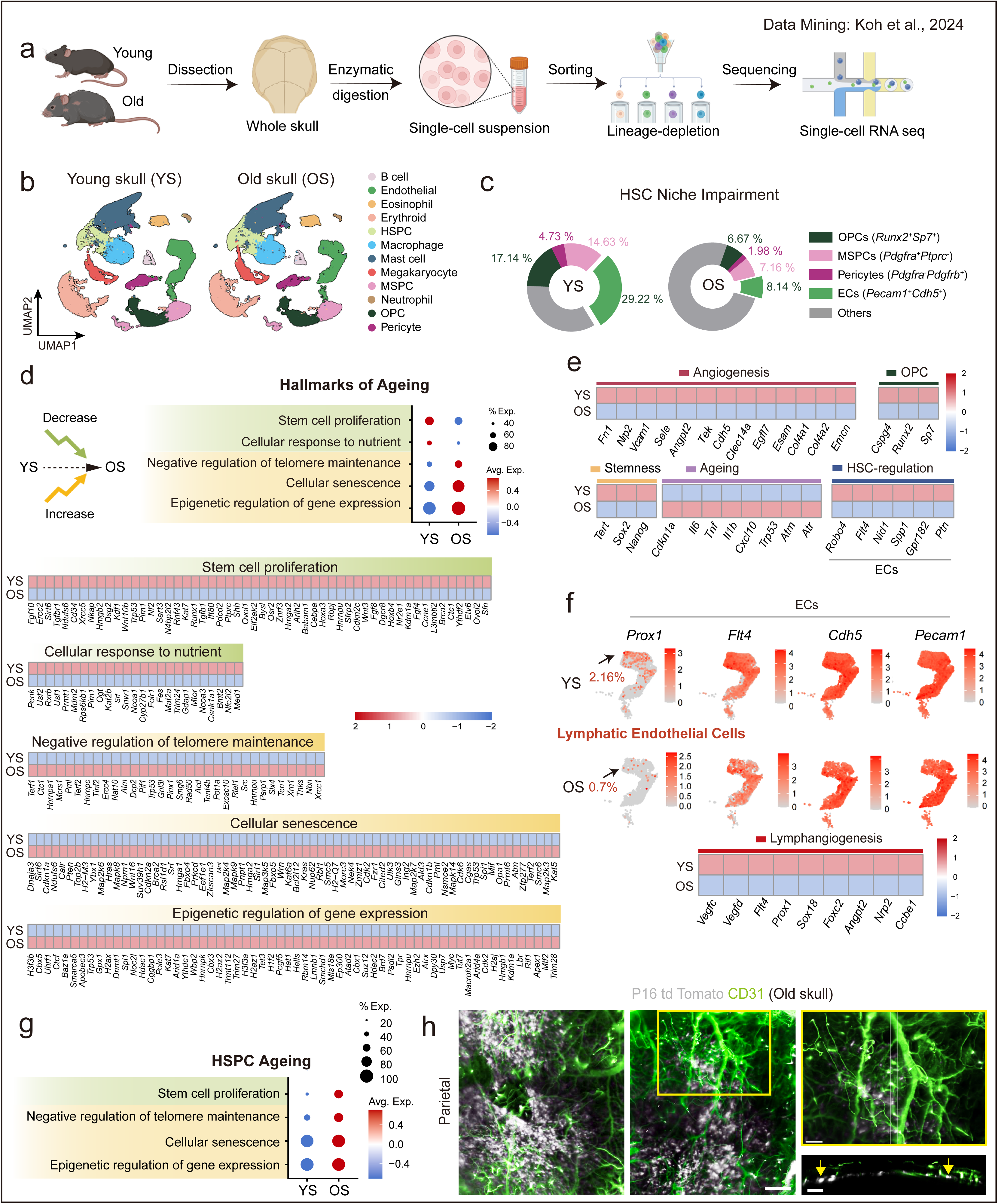
Unbiased re-analysis of DOI: 10.1038/s41586-024-08163-9 reveals hallmarks of ageing in murine skull HSPCs and niche populations. a, Schematic of experimental work-flow used in DOI: 10.1038/s41586-024-08163-9 to obtain scRNA-seq data (GSE275179) from skulls of young and old mice. b, UMAP projections showing distribution of various cell populations in young skulls (YS; GSM8472479-GSM8472481, GSM8472486) and old skulls (OS; GSM8472469, GSM8472482, GSM8472484). c, Pie charts illustrating the proportions of osteoprogenitors (OPCs), mesenchymal stem and progenitor cells (MSPCs), pericytes, and endothelial cells (ECs) in YS and OS. d, Transcriptomic signature showing GO terms derived from the Molecular Signatures Database (MSigDB) for YS and OS, with heatmaps showing log-normalized expression of related genes (z-scored per gene across groups), in the context of hallmarks of ageing. e, Colour-coded heatmaps displaying log-normalized expression of selected genes associated with angiogenesis, osteoprogenitors, stemness, ageing, and HSC regulation in YS and OS. f, *Prox1*, *Flt4*, *Cdh5*, and *Pecam1* expression patterns in the EC cluster with lymphatic ECs defined as *Cdh5⁺ Pecam1⁺ Ptprc⁻ Flt4⁺ Prox1⁺* cells. Colour-coded heatmap showing log-normalized expression of selected genes associated with lymphangiogenesis in YS and OS. g, Transcriptomic signature scoring of GO terms derived from MSigDB for HSPCs identified in YS and OS in the context of hallmarks of ageing. h, Representative 3D images of parietal region of skull from a 100-week-old P16 td Tomato mouse showing blood vessels with CD31 (green) and in vivo p16-expressing senescent cells (white). The white dotted lines indicate the region of the 80-µm thick orthogonal view shown. Arrows indicate the p16 expressing endothelial cells. Scale bars: 500 μm (low magnification); 300 μm (high magnification and orthogonal view).

Next, we performed unbiased proteomic profiling of young mice (10 weeks) versus old mice (79 weeks) skulls to independently validate the transcriptional findings (Fig. 2a, b). Consistent with the single-cell RNA-seq analysis, old skull bone and bone marrow exhibited an impairment of niche-supportive programs, reflected by downregulation of proteins associated with mesenchymal stem cell and osteoprogenitor cell identity. Global pathway analyses revealed marked enrichment for inflammation and immune activation in old skulls, including increased abundance of proteins linked to immune response and antigen processing and presentation (Fig. 2c). In parallel, mitochondrial proteins, such as components of complex I (NADH dehydrogenase iron sulphur- and flavo-proteins) and V (ATP synthase subunits) of the electron transport chain, proteins involved in mitochondrial proteostasis (LONP1) and oxidative homeostasis (MSBR3 and ROMO1), were significantly reduced (Fig. 2d), suggesting mitochondrial dysregulation and metabolic decline. Notably, there were greater levels of DNA damage-associated proteins including proteins involved in nucleotide-excision repair (RAD23A/B), DNA-protein crosslink proteolysis (FAM111A), and DNA interstrand crosslink repair (NEIL3) in old skulls (Fig. 2d). We also detected increased levels of proteins that respond to DNA-protein or DNA-intrastrand crosslinks, such as the MCM complex, replicative DNA polymerases, and REV3L, the catalytic subunit of translesion DNA polymerase zeta, which bypasses DNA lesions after DNA-protein or DNA-intrastrand crosslink resolution. In addition, we found higher expression of proteins linked to DNA damage-induced HLA presentation (TAP1, TAP2) (Fig. 2d). Alongside the broad upregulation of DNA damage-associated proteins, the proteomic data also indicate that OS cells experience DNA replication stress, a hallmark of sporadic cancer, as evidenced by elevated levels of replication stress markers such as ATR, CHEK1, and RPA (Fig. 2d). Pathways critical for vascular and skeletal integrity were also impaired: proteins associated with angiogenesis including VEGF receptor NRP2 and Cadherin-5, osteogenesis, such as osteoblast-specific transcription factors (RUNX2 and OSX), extracellular matrix components with osteogenic stimuli were sharply diminished, whereas those linked to inflammation and cellular senescence were strongly elevated (Fig. 2d). Not only this, proteins in CD200-CD200R1 axis known for its role in limiting inflammation were also found to decrease in OS (Fig. 2d). Together, these proteomic signatures strongly reinforce the single-cell RNA-seq findings, demonstrating that the aged skull bone marrow (BM) niche undergoes coordinated cellular and molecular deterioration across multiple layers of regulation. These changes reflect the activation of canonical hallmarks of ageing, including DNA damage, senescence, altered metabolism, and inflammation. These cellular reductions were accompanied by transcriptional programs indicative of canonical ageing hallmarks, including cellular senescence, impaired telomere maintenance, and reduced stem cell proliferation.

**Fig. 2:**
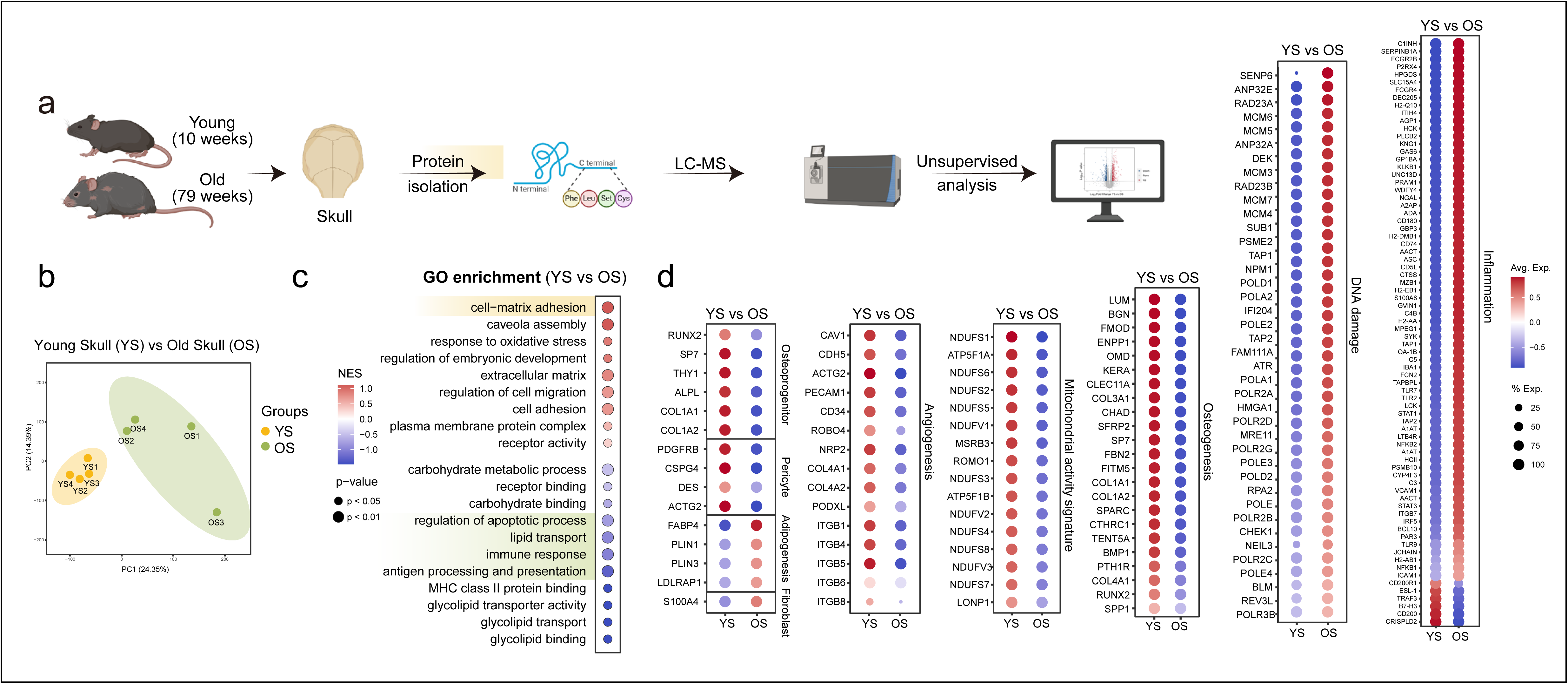
Unbiased proteomics reveals age-dependent impairment of skull bone and bone marrow, highlighting the emergence of ageing hallmarks. a, Schematic of proteomic profiling of skulls from young and old mice. b, Principal component analysis (PCA) of proteomic profiles from young (yellow) and old (green) skulls. (n=4 per age group) c, GO enrichment analysis of proteomic profiles of YS and OS identifies age-associated alterations in biological processes. d, Dot plots showing differential expression of proteins linked to osteoprogenitors pericytes, adipogenesis, fibroblasts, angiogenesis, mitochondrial activity, osteogenesis, DNA damage, and inflammation in young skulls (YS) versus old skulls (OS). Colour and size of the dot indicate the log expression value of the protein and the fraction of animals expressing the protein respectively.

It is important to note that the relative abundances of vascular cells in scRNA-seq datasets are influenced by technical factors such as tissue dissociation and sample processing^21–24^. In both skulls and femurs, endothelial cells, including sinusoidal endothelial cells which are *Vcam1^+^*, are consistently underrepresented, with the effect being most pronounced in the skulls. This likely reflects the heightened fragility and senescent state of ageing endothelial populations, particularly sinusoidal endothelial cells (*Vcam1^+^*), which are preferentially lost during single cell sequencing workflows^25^. Immunofluorescence analyses of bone marrow regions in young versus old skulls provided spatial context for the observed molecular changes. Fibroblast marker FSP1 revealed a notable expansion of fibroblasts in old skull marrow (Extended Data Fig. 1c), indicating remodelling of the mesenchymal compartment^26^. Markers of DNA damage and senescence, γ-H2AX and p16, were elevated in the bone marrow of aged skulls (Extended Data Fig. 1d, e), reflecting replication stress. CD3 and CD8 immunostaining analyses on bone marrow showed increased accumulation of immune cells, suggesting an enhanced inflammatory response (Extended Data Fig. 2a). BODIPY and Perilipin staining demonstrated elevated adipogenesis in bone marrow regions from old skulls (Extended Data Fig. 2b). Levels of the angiogenic factor VEGFA and the proliferation marker Ki67 were reduced, indicating compromised angiogenic signalling and reduced proliferation, while immunostaining for the osteoprogenitor marker OSX was diminished, reflecting impaired osteogenic potential (Extended Data Fig. 2c, d). Collectively, these results indicate that the skull bone marrow and the bone marrow niche undergo substantial structural and functional decline with age, affecting both stromal and hematopoietic compartments.

### Skull Bone Marrow Maintains Widespread Vasculature in Young Mice

Contrary to the report^1^ describing only restricted vascularized areas in young skull bone marrow, with supposed expansion during ageing, our analyses reveal a fundamentally different vascular organization. Endomucin immunofluorescence in their study^1^ is localized to nuclei (Extended Data Fig. 3), which is inconsistent with its established role as a transmembrane endothelial cell surface marker^19,20^. Our imaging of skull showed that Endomucin immunofluorescence localise to cell surface (Fig. 3a).

**Fig. 3:**
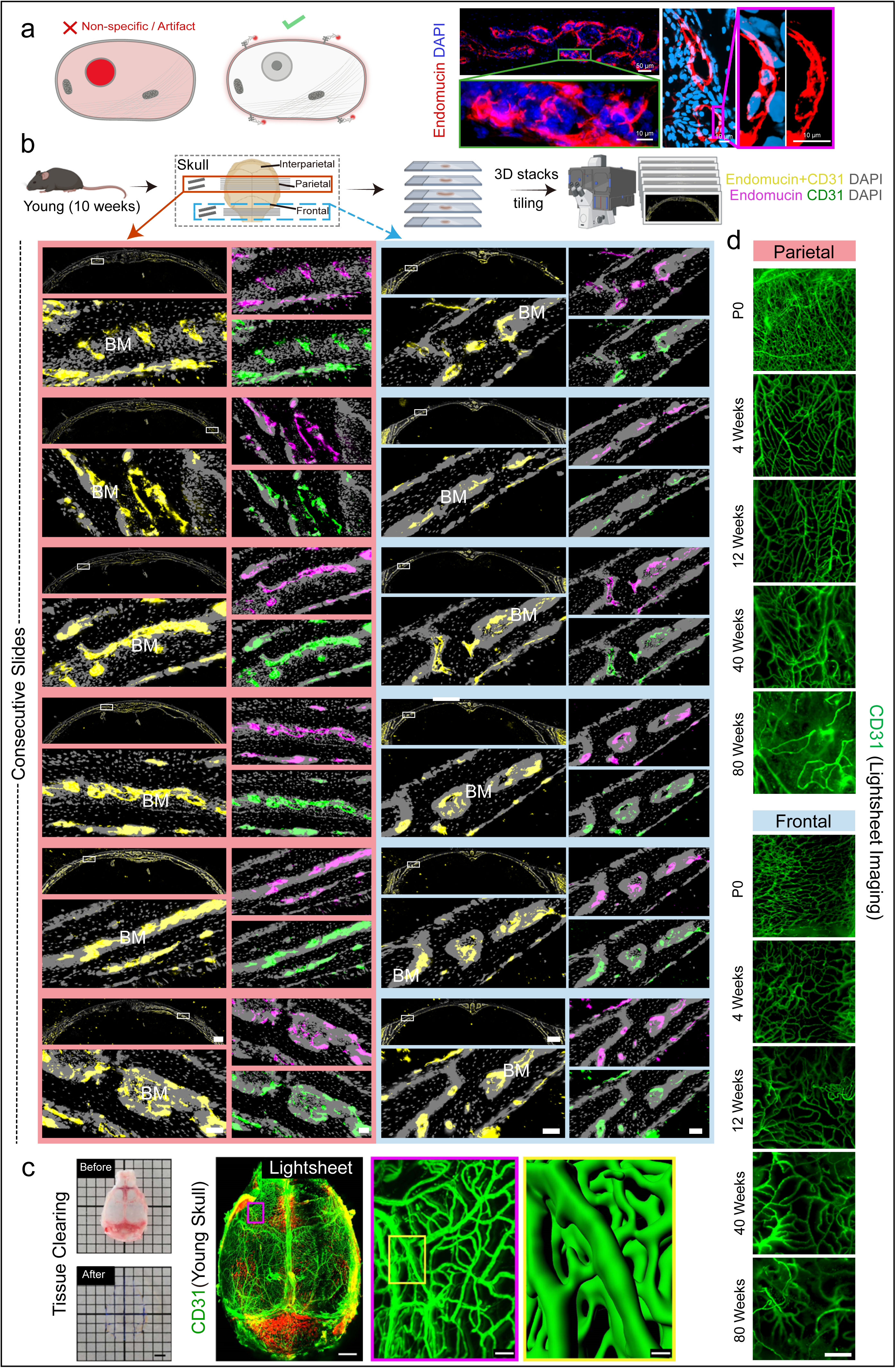
Parietal and frontal regions of murine young skulls harbour a continuous vascular network. a, Schematic illustrating non-specific immunofluorescence versus true membrane localization. Representative 3D immunofluorescence images of Endomucin (red) with DAPI (blue) in the parietal region skull from a young mouse, confirming specific endothelial cell surface localization. Scale bars: 50 μm (overview images); 10 μm (high-magnification images). b, Experimental scheme for thick-sectioning of parietal and frontal regions of skulls from young (10 weeks) mice, followed by 3D imaging and vascular analysis. Representative maximum-intensity projections of consecutive sections from the parietal and frontal regions of skulls from young mice, labelled with nuclear stain DAPI (grey) and immunostained for Endomucin (magenta) and CD31 (green), showing Endomucin+CD31 cells in yellow. BM: bone marrow; Scale bars: 500 μm (overview images); 50 μm (all other images). c, Young skull prior to and after tissue clearing. 3D images of the whole skull showing CD31 expression pattern. Scale bars: 500 μm (whole skull); 50 μm (middle); 10 μm (right). d, Representative images of skull marrow from frontal (top) and parietal (bottom) regions of postnatal day 0, 4 weeks, 12 weeks, 40 weeks and 80 weeks old mice showing CD31^+^ vessels (green). Scale bar: 300 μm

Using immunohistochemistry of thick sagittal sections and light sheet imaging of cleared whole skulls, we consistently observed a continuous, dense vascular network throughout the skull frontal, parietal and interparietal regions in young mice (Fig. 3b, c; Extended Data Fig. 4). Serial analyses of parietal sections confirmed widespread vascular presence, directly contradicting the proposed existence of avascular domains within the parietal bone (Fig. 3b). Light sheet imaging further substantiated these findings, capturing an extensive vascular network spanning the entirety of young and old mouse skulls (Fig. 3c, d; Movie 1; Extended Data Fig. 5a).

The discrepancy between our findings and the earlier report^1^ is likely attributable to methodological differences. The previous study used uncleared whole-mount skulls, limited imaging depth with confocal microscopy, lower resolution, incomplete coverage of tissue and bone marrow, and sampling strategies that may have introduced artifacts^1^. Our whole-mount preparations consistently revealed DAPI-stained nuclei throughout the skull marrow, whereas their skull imaging shows regions absent or weak nuclear signals (Extended Data Fig. 5b). Collectively, our results demonstrate that the skull bone marrow in young mice is rich and continuously vascularized. These findings challenge the proposed model of limited vascularization and non-vascularized in young skulls and highlight the importance of high-resolution, whole-skull imaging across multiple regions. By providing this comprehensive view, our study establishes a robust baseline for the vascular architecture of the skull bone marrow and emphasizes the need for careful methodological validation to accurately interpret tissue organization.

### Bone Marrow Expansion with Age Is Not Unique to the Skull

The previous study^1^ suggested that bone marrow expansion with ageing is a unique feature of the skull, based on limited comparisons with femurs. However, reanalysis of their data shows that aged long bones also exhibit increased frequencies of CD45⁺ hematopoietic cells and HSPCs (Fig. 4a). Single-cell RNA-seq analysis of femurs reveals hallmarks of ageing similar to those observed in the skull, including increased cellular senescence and loss of angiogenic programs (Fig. 4b-d). Our structural analyses of long bones show significant increases in overall length and dimensions with age (Fig. 4e). Micro-CT analyses further demonstrate expansion of the bone marrow cavity along with the well-documented loss of trabecular structures in the metaphysis, a hallmark of skeletal ageing (Fig. 4f-h; Movie 2). The combination of dimensional enlargement and trabecular bone loss creates additional space that is progressively occupied by hematopoietic cells. Together, these findings indicate that the marrow expansion previously attributed to the skull likely reflects a misinterpretation from skull-focused analyses without proper cross-skeletal comparison. Collectively, this challenges the notion of skull-specific marrow expansion and underscores the importance of rigorous anatomical controls when studying bone marrow dynamics across the skeleton.

**Fig. 4:**
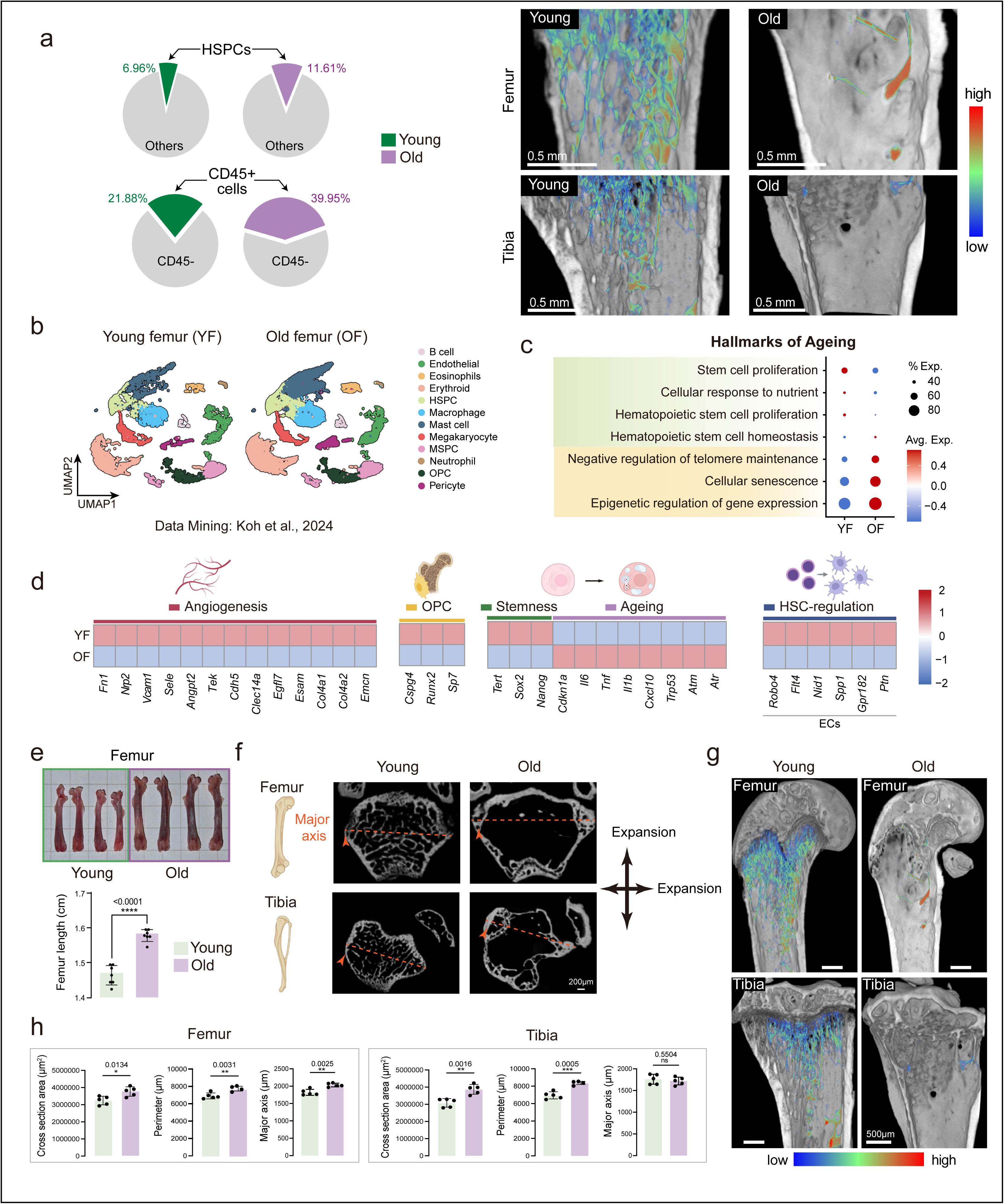
Long bones undergo transcriptional, structural and cellular remodelling with ageing leading to bone marrow expansion. a, Pie charts showing the proportion of HSPCs and CD45^+^ cells identified in young (YF; GSM8472485) versus old femurs (OF; GSM8472468 & GSM8472483) from re-analysis of publicly available DOI: 10.1038/s41586-024-08163-9 single cell RNA-seq data set GSE275179. Representative micro-CT images of femur and tibia from young and old mice, highlighting trabecular bone loss in old mice. b, UMAP projections illustrating the distribution of various cell populations in young femurs (YF) and old femurs (OF), colour coded by identified cell types. c, Transcriptomic signature scoring of GO terms derived from the Molecular Signatures Database (MSigDB) for YF and OF, in the context of hallmarks of ageing. d, Colour-coded heatmaps showing the log-normalized expression of selected genes associated with angiogenesis, OPCs/osteoprogenitors, stemness, senescence, and HSC regulation in YF and OF. e, Representative photos of femurs from young and old mice, with corresponding quantification of femur length (*n* = 7 animals per age group). f, Representative 2D micro-CT images (top views) of femur and tibia from young and old mice. g, Representative 3D micro-CT images of femur and tibia from young and old mice, highlighting trabecular bone in colour scale. h, Quantification of 2D femur images from young and old mice showing cross-sectional area, perimeter, and major axis (*n* = 5 animals per age group). Data are presented as mean ± s.d. Statistical tests: (e) Welch’s t-test; (h) two-tailed unpaired Student’s t-test.

### Compared with the skull, vertebrae exhibit relative resistance to the hallmarks of ageing

Vertebrae are the principal source of hematopoietic cells and the standard site for bone marrow harvest in clinical transplantation. To directly compare ageing-associated changes across bone marrow sites, we performed unbiased proteomic analyses of skull and vertebrae from the same old mice (Fig. 5a, b). This analysis revealed striking divergence: whereas vertebrae appeared relatively protected, old skull bone marrow displayed pronounced upregulation of inflammatory proteins, immune activation, DNA damage and replication stress pathways, as well as increased peptidase activity and tRNA methylation, processes previously implicated in age-associated proteostasis imbalance (Fig. 5c, d).

**Fig. 5:**
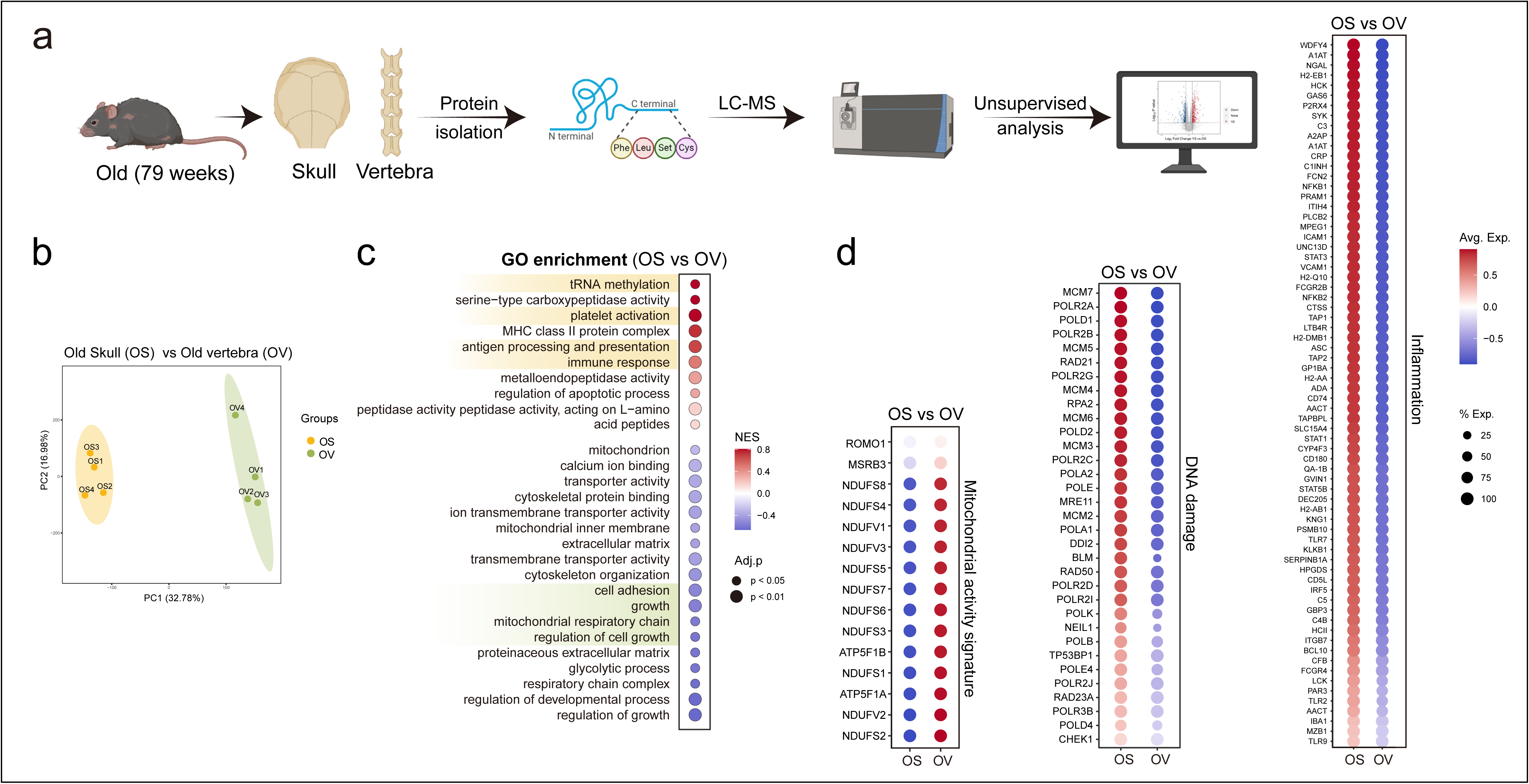
Proteomic profiling reveals vertebrae are relatively protected from ageing hallmarks compared to skull. a, Schematic overview of proteomic profiling of skulls and vertebrae from old mice to analyse bone-specific protein signatures. b, Principal component analysis (PCA) of proteomic profiles from old skull (OS; yellow) and vertebrae (OV; green) (*n* = 4 mice per age group). c, GO enrichment analysis of proteomic profiles of old skull samples (OS) versus old vertebra samples (OV). d, Dot plots illustrating differential expression of proteins linked to mitochondrial activity, DNA damage, and inflammation. Colour and size of the dot indicates the log expression value of the protein and the fraction of mice expressing the protein respectively.

We did not directly compare resident stromal, endothelial, or bone cells; the distinct developmental origins of intramembranous versus endochondral ossification, together with the structural differences between flat and long bones, likely contribute to these divergent ageing trajectories. Proteomic profiling revealed that old vertebrae, relative to skull, exhibited upregulation of pathways related to cell adhesion and migration (Fig. 5c), while mitochondrial proteins associated with oxidative phosphorylation and regulation of oxidative state were downregulated in old skulls, indicating mitochondrial dysregulation and altered metabolism (Fig. 5d).

Imaging of neutral lipids and the lipid droplet marker protein Perilipin displayed low abundance of adipocytes (Extended Data Fig. 2b; Extended Data Fig. 6). These findings collectively demonstrate that the skull bone and its bone marrow undergo profound ageing-related degeneration, whereas vertebrae are relatively protected from hallmarks of ageing (Fig. 6). This differential susceptibility may reflect both developmental origin and local microenvironmental factors, with potential implications for skeletal ageing and clinical hematopoietic interventions.

**Fig. 6:**
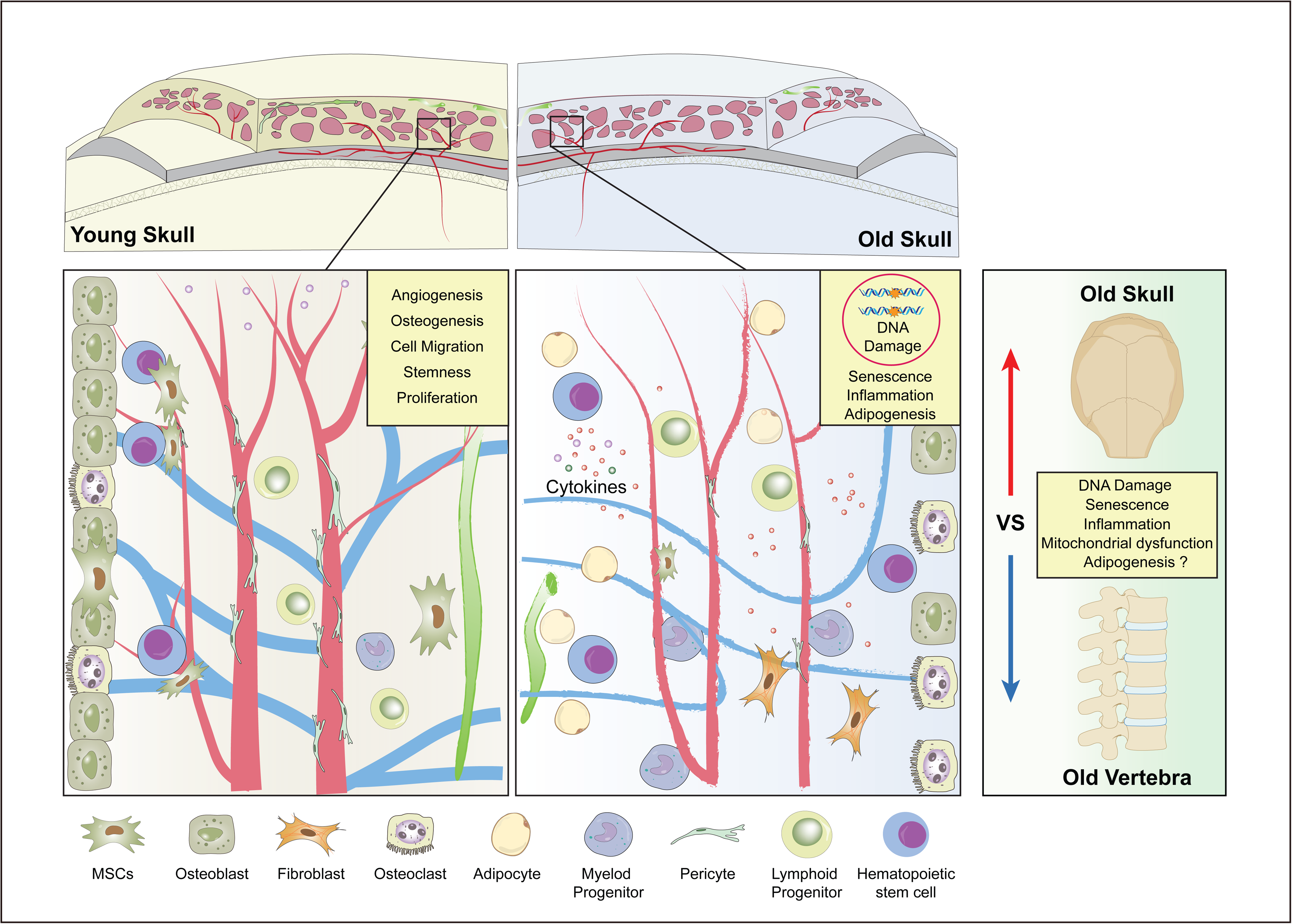
Schematic showing degeneration and impaired resilience of skull bones.

## Discussion

Skull bone is shown to undergo age-associated deterioration^4,5,27^ and skull bone and bone marrow altered in age-associated neurological diseases^28^. Aging leads to bone loss, bone thinning and induces structural alterations^29,30^. The notion that the skull bone marrow constitutes a uniquely expanding and resilient reservoir during ageing is not supported by evidence. Across multiple experimental and computational approaches, we find the opposite: the skull marrow niche undergoes profound and multifaceted deterioration. Mesenchymal progenitors are depleted, angiogenic and lymphatic programs decline, adipocytes accumulate, and hallmarks of genomic and cellular ageing—DNA damage, replication stress, senescence, and inflammation—dominate the old skull bone and bone marrow. These changes are not subtle. They are striking, reproducible, and consistent across independent analyses. Many of the cellular changes and hallmarks of ageing demonstrated by such unbiased analyses has been reported as a feature of bone and haematopoietic ageing^3,14,15,31–34^. Moreover, high-resolution imaging refutes the idea of “avascular zones” in young skulls: the parietal region is well vascularized, directly contradicting one of the central assertions of the prior report^1^. The skull is not preserved during ageing—it is among the most vulnerable sites of marrow ageing.

We note major methodological flaws that undermine the reliability of their data. Imaging uncleared whole-mount skulls with confocal microscopy lacks the necessary depth to resolve bona fide marrow structures, and the Endomucin staining shows aberrant nuclear localization that is inconsistent with a cell-surface endothelial marker. No justification or discussion is provided for this unexpected nuclear signal. Moreover, the data from whole-skull imaging and single-cell sequencing are presented as “bone marrow” without accounting for contributions from bone and cortical regions.

The GEO datasets raise further concerns: experimental groups are not clearly linked to figures and panels, sample labelling is inconsistent, and HSPC isolation protocols differ from those reported^1^. Notably, sample names were altered in the GEO datasets many months after the publication and initial deposition, without any justification provided for these changes.

Collectively, these issues compromise data transparency and reproducibility, casting serious doubt on the claim that skull marrow represents a uniquely resilient haematopoietic reservoir. Similarly, cellular analysis of adipocytes in skull versus femur requires careful consideration, as differences in bone type, marrow compartments, structural organization, and ossification patterns make direct comparisons problematic.

Our analyses establish an accurate framework: ageing does not spare the skull marrow but drives its accelerated degeneration relative to vertebrae, the principal haematopoietic reservoir. This compartment-specific vulnerability has important implications. Skull marrow connects directly with meningeal vessels and lymphatics, suggesting that its accelerated decline may contribute to impaired neuro-immune surveillance in older individuals. Such changes could influence stroke recovery, neurodegenerative disease progression, and age-associated neuroinflammation.

Clinically, the relative preservation of vertebral marrow provides a reassuring explanation for why many elderly patients retain hematopoietic reserve sufficient for transplantation, while others experience premature immune frailty. It highlights the need for cross-skeletal validation: a single aspirate or biopsy may not reliably represent the ageing marrow.

Finally, our identification of senescence, replication stress, mitochondrial dysfunction, and TGFβ signalling as key drivers of skull marrow ageing points to actionable pathways. These findings open new opportunities for therapeutic intervention and reframe the ageing marrow not as a uniform process, but as a compartmentalized and clinically relevant landscape of resilience and vulnerability.

## Acknowledgments

The author(s) declare that financial support was received for the research. A. P. K. is supported by Ministry of Education (MOE) Singapore: Academic Research Funds (#024983-00001), European Research Council (StG: metaNiche, 805201), European Union’s Horizon 2020 (No 857524). S. K. R. acknowledges funding support from the Ministry of Education (MOE), Singapore: SUG (024482-00001) and Academic Research Funds Tier 1 (#024712-00001) and European Union’s Horizon 2020 (No 857524). W. L. G. is supported by National Institute of Dental and Craniofacial Research (NIH-NIDCR:1R01DE027957; NIH-NIDCR: 1F31DE033910). K.R. received support from the Medical Research Council programme (Grant Ref. MR/X006409/1), Breast Cancer Now (Grant Ref. 2022.11PR1570), Ministry of Education-Strat-Up Grant, Singapore (023917-00001), Toh Kian Chui Distinguished Professorship Award/LKC Medicine, Singapore. C.C. is supported by the Ministry of Education Academic Research Fund (RG34/25), Singapore. N. K. V. acknowledges funding support, in part, by the National Research Foundation Singapore under its Open Fund - Individual Research Grant administered by the Singapore Ministry of Health’s National Medical Research Council (#MOH-001892). M.V.R. acknowledges support by the National Institute on Aging (R01AG073349) and National Institute of Arthritis and Musculoskeletal and Skin Diseases (R01AR055655 and R01AR082460). J. C. is supported by the National Nature Science Foundations of China (Nos. 82422021, 82270961), and Sichuan Science and Technology Program (No. 2023JDRC0018). R. D. is supported by Personalised Cardiometabolic Risk Management (predict-2-prevent) and the vascular research initiative supported by National Heathcare Group and Lee Kong Chian School of Medicine. A.B. received funded DFG-SFB/TRR369 DIONE-501752319. L.Y. received funding from MOE-SUG (022976-00001) and NRF-CRP (NRF-CRP24-2020-0004). A. C. is supported by R01 AG082681 grant. S. J. and M.S.K. are supported by the Nanyang Technological University Research Scholarship (Reg. No. 200604393R). Y.Y. is supported by National Nature Science Foundations of China (No. 824B2026).

## Methods

### Mice

C57BL/6J strain was used as wild-type mice in all experiments. Mice aged 9-11 weeks were used in young groups and 79-82 weeks were used in old groups unless stated otherwise were designated as young and old groups, respectively. Both male and female mice were used in the study. For skull bone analyses the dura mater was carefully removed from the skull. Cdh5-2A-CreERT2 (NM-KI-200173) and Rosa26-LSL-tdTomato (NM-KI-225042) strains were supplied by Shanghai Southern Model Biotechnology Co., Ltd. Experiments involving animals were conducted in accordance with local ethical standards and obtained approval from Nanyang Technological University, Singapore; Imperial College London, UK; University of Oxford, UK; Johns Hopkins University, USA; Sichuan University China and Mayo Clinic USA.

### Single cell RNA-seq analysis

Single-cell RNA-seq datasets were retrieved from the Gene Expression Omnibus. Raw UMI count matrices in 10X Genomics format were imported into R (v4.4.1) with Seurat (v5.2.1) Read10X and converted to individual Seurat objects via CreateSeuratObject. Objects were merged (merge) while retaining sample-level metadata to preserve experimental group identities.

Quality control (QC) metrics were computed per cell, including the number of detected genes (nFeature_RNA), total UMI counts (nCount_RNA), and the fraction of mitochondrial transcripts (using PercentageFeatureSet with the patterns ^mt-for mouse or ^MT-for human). Cells were retained if 500–6,000 genes were detected and mitochondrial RNA accounted for <25% of total UMIs. QC distributions were visualised with VlnPlot.

Counts were normalised with NormalizeData (method = “LogNormalize”) and scaled using ScaleData. Highly variable genes (HVGs, n = 2,000) were selected with FindVariableFeatures (method = “vst”). Principal component analysis (PCA) was performed with RunPCA, and the number of informative components was guided by elbow plots (ElbowPlot); 50 PCs were retained for downstream steps. To account for between-sample effects, batch correction was performed with Harmony (v1.2.3) using RunHarmony with sample identity as the covariate. Non-linear dimensionality reduction was performed with UMAP (RunUMAP) on the Harmony-corrected embeddings. Graph-based clustering was carried out with FindNeighbors and FindClusters on the corrected dimensions across a range of resolutions, and cluster stability across resolutions was assessed with clustree (v0.5.1). Heatmaps were generated from Seurat log-normalized data. For each gene, group means were computed with AverageExpression (assay: RNA) and then z-scored per gene across groups; matrices were visualized with heatmap. GO term signatures were retrieved from MSigDB, and per-cell scores were computed using AddModuleScore function from Seurat.

Cell types were assigned by manual inspection of canonical markers, visualised with FeaturePlot, VlnPlot, and DotPlot. Relative cluster proportions across conditions were computed using reshape2 (v1.4.4). Figures were assembled with ggplot2 (v3.5.1) and patchwork (v1.3.0).

Key resources: Seurat v5.2.1^35^; Harmony v1.2.3^36^; clustree v0.5.1^37^.

### Proteomics

#### Sample collection and storage for proteomics

Young (10-week-old) and old (79-week-old) wildtype C57BL/6J mice were euthanized following approved institutional procedures. Skull and vertebrae were immediately dissected kept on ice, then briefly rinsed with cold PBS to remove blood and debris. Tissues were labelled, snap-frozen in liquid nitrogen for 5 minutes, and stored at −80 °C until proteomic analysis. No chemical fixation or decalcification was performed, ensuring preservation of the native protein composition.

### Protein extraction

Proteins were extracted using the Novogene standard operating procedure. Briefly, frozen tissues were pulverized in liquid nitrogen and lysed in SDT buffer (100 mM NaCl; DTT 1:100, v/v). Lysates were sonicated on ice, heat-denatured (95 °C), cleared (12,000 g, 4 °C), and alkylated with iodoacetamide (room temperature, dark). Proteins were precipitated with pre-chilled acetone (1:4, v/v, −20 °C), washed, air-dried, and re-dissolved. Aliquots were normalized to 100 µL and digested with trypsin in two steps (4 h, then overnight at 37 °C). Peptides were acidified (pH < 3), clarified, desalted on C18, dried, and reconstituted; ∼200 ng were injected. iRT standards and Bradford assays were used as needed for QC and loading.

Peptides were separated on a Vanquish™ Neo UHPLC with a C18 trap–analytical setup using water/0.1% formic acid (A) and acetonitrile/0.1% formic acid (B) under a short gradient, followed by a high-organic wash. MS data were acquired on a Thermo Orbitrap Astral (Easy-Spray source; 2.0 kV; 290 °C) in DIA mode. MS1 scans spanned m/z 380–980 at 240k resolution; DIA employed 2 Th windows across the MS1 range (NCE 25). MS2 spectra were recorded at 80k resolution with standard injection times. Raw data were saved as *.raw files.

### Identification and quantification

DIA files were processed in DIA-NN. The search used a reference protein database matching the species. Precursor tolerance was 10 ppm; fragment tolerance was 0.02 Da. Carbamidomethyl-C was fixed. Oxidation (M) and protein N-terminal acetylation (with Met loss options) were variable. Up to two missed cleavages were allowed. PSMs, peptides, and proteins met an FDR of 1% or lower. iRT was used for retention-time correction. Protein abundances were compared between groups. Multiple testing control was applied. Differential proteins met two rules: |log2 fold-change| ≥ log2(1.5) and P < 0.05. Functional analysis used standard enrichment on GO/KEGG/Reactome with all identified proteins as background.

### Micro-computed tomography (Micro-CT)

Female C57BL/6 J mice (Charles River, strain 632) were housed in groups housed (2–5 mice/cage) under 7 a.m.-7 p.m. lighting with food and water ad libitum. Tissues fixed in 4% paraformaldehyde were scanned using a Skyscan 1276 micro-CT system (Bruker) with the following acquisition parameters: filter: Al 0.5 mm, source voltage = 70 kV, source current = 200 µA, pixel size = 4 µm, exposure time = 1114 ms, rotation step = 0.2°. Image reconstruction was performed using NRecon software (Bruker), and 3D quantitative analysis of bone parameters was conducted using CTAn software (Bruker, version 1.20.8.0) and Fiji (ImageJ, version 1.54p). Calvaria ROI: anterior area of the parietal bone adjacent to Bregma, 2 mm wide. Femur ROI: distal area of trabecular bone adjacent to the growth plate, 800 µm long. Tibia ROI: proximal area of trabecular bone adjacent to the growth plate, 800 µm long.

### Tamoxifen treatment

For oral gavage, tamoxifen (Sigma-Aldrich, T5648) was initially dissolved in 100% ethanol and subsequently diluted in corn oil to reach a final concentration of 5 mg/ml. To induce Cre activity of Cdh5-2A-CreERT2 X Rosa26-td-Tomato transgenic mice, tamoxifen was administered orally at a dose of 50 mg/kg for five consecutive days. The mice were euthanized and examined 2 weeks after the final tamoxifen treatment.

### Skull fixation, and decalcification for light sheet imaging

Light sheet imaging was performed as previously described^5,38^. After dissection and removal of surrounding tissues, skull samples were fixed in 4% paraformaldehyde (PFA; Sigma-Aldrich, P6148) for 2.5 h. Dura mater was also carefully removed. Decalcification was carried out in 0.5 M EDTA (Sigma-Aldrich, E5134) (pH 7.4) at room temperature for 24 h. Bones were rinsed in PBS and then dehydrated using an ice-cold methanol (Sigma-Aldrich, 179337) series: 50% and 80% for 30 min each, then 100% for 1 h, with the methanol replaced twice every 20 min. After bleaching in 5% H₂O₂ (Sigma-Aldrich, HX0636) for 1 hour, tissues were rehydrated with a decreasing methanol gradient and rinsed in PBS.

For antigen retrieval and permeabilization, tissues were incubated at 4 °C for 5 h in a solution containing 25% urea (Sigma, U5128), 15% Triton X-100 (Sigma, T8787), and 15% glycerol (Sigma, G7893), followed by 0.2% collagenase (Merck, 10103578001) digestion at 37 °C for 30 minutes. Samples were washed with 2% FBS (Sigma-Aldrich, F7524) in PBS and then blocked in 10% donkey serum (Abcam, ab7475), 10% DMSO (Sigma-Aldrich, D5879), and 0.5% Triton X-100 in PBS for 20 min at 37 °C. Primary antibodies (1:300) were diluted in buffer containing 0.5% Triton X-100, 2% donkey serum, and 10% DMSO in PBS, and samples were incubated overnight. Subsequent washes were performed in PBS containing 2% donkey serum and 0.5% Triton X-100 at 37 °C with shaking (70 rpm) for 3 h, replacing the wash buffer every 15 min during the first hour and every 30 min thereafter. Fluorophore-conjugated secondary antibodies (1:500 dilution) were applied for 6–8 h at 37 °C, followed by the same wash procedure. Immunolabeled skulls were dehydrated through a graded isopropanol (Sigma-Aldrich, 190764) series (30%, 50%, 80% and 100%) and cleared in a solution of 80% ECi (Sigma-Aldrich, 112372) and 20% PEGM (Sigma-Aldrich, 447943). The Zeiss Lightsheet 7 microscope with a 5X EC Plan-Neofluar objective was used for image acquisition. Samples were mounted with cyanoacrylate in ECi, and image datasets were processed using Imaris software (version 10.2.0).

### Immunostaining on skull sections

Bone samples were prepared, sectioned and immunostained as described previously^9^. Fresh samples were fixed in 4% PFA for 4 h at 4 °C. The skull specimens were then demineralized in 0.5 M EDTA (pH 7.4) for at least 24 hours. Tissues underwent overnight cryoprotection at 4 °C in 20% sucrose and 2% polyvinylpyrrolidone (PVP). Samples were then embedded in a freezing solution consisting of 20% sucrose, 8% gelatin, and 2% PVP. Thick sections (20–80 μm) were prepared on a cryostat using low-profile blades and air-dried for 5 h before being frozen for subsequent immunostaining. Following thawing, sections were air-dried for 15 min, rehydrated for 5 min, permeabilized with 0.3% Triton X-100 for 10 min, and blocked at room temperature with 5% donkey serum. Primary antibodies at a 1:150 dilution were applied to sections in blocking solution for 4 h at room temperature, followed by secondary labelling with Alexa Fluor-conjugated antibodies (1:300) for 90 min. For nuclear staining, DAPI was applied, and sections were mounted in Fluoromount-G (Invitrogen, 00-4958-02). Imaging was conducted using three different microscopes: a Dragonfly 200 spinning disk confocal microscope (Andor, Oxford), a Zeiss LSM880 laser scanning confocal microscope, and a Leica DM6 B Thunder Microscope. Immunofluorescent bone sections were imaged using the Leica DM6 B Thunder Microscope (LAS X v3.7.4.23463), and tile scans were stitched and refined with Thunder Computational Clearing to reduce background haze and enhance contrast. All three-dimensional immunofluorescent datasets obtained from these microscopes were subsequently processed and reconstructed using Imaris software (version 10.2.0). All antibodies employed are detailed in Tables S1 and S2.

### BODIPY staining

For neutral lipid staining in thick bone sections, 80-μm cryosections of different bones were incubated with 0.5 μM BODIPY 558/568 (Invitrogen, D3835) for 30 min at room temperature. Sections were subsequently washed three times with PBS at 5-min intervals and then incubated with DAPI (1:1000 dilution) for 30 min. Following additional PBS washes, the stained samples were carefully mounted with glass coverslips using Fluoromount-G (Invitrogen, 00-4958-02).

### Imaging quantifications

To quantify the coverage percentage of Perilipin, VEGFA, Ki67, OSX, CD3, CD8, FSP1, and γ-H2AX, ImageJ (version 1.53t) was employed. Regions of interest were first defined, and the positively stained areas were identified and measured. The proportion of coverage was determined as the ratio of stained area to the total area within the selected region.

### Statistical analysis

Multiple and independent experiments were performed to validate the reproducibility of findings. Sample sizes were not pre-determined based on statistical power calculations. Mice were allocated to experiments randomly, and samples were processed in an arbitrary order. No blinding was performed. No animals were excluded from analysis. All statistical analyses were conducted with GraphPad Prism software (version 9.2). All data are expressed as mean ± s.d. Two-tailed Student’s unpaired t-test or Welch’s t-test was applied for the analysis of the statistical significance of differences. P < 0.05 was considered significant. ns indicates not significant; *: P < 0.05; **: P <0.01; ***: P < 0.001; ****: P <0.0001.

### Data and Code availability

Analysis codes for the single-cell sequencing and proteomics workflows are provided in the Supplementary Files. The proteomic raw data generated in this study have been deposited in the iProX under accession number IPX0013567000, and will be publicly available upon publication of this article.

---

**Extended Data Fig. 1: Ageing drives degeneration of the skull bone marrow and the bone marrow microenvironment**

a, Pie charts showing the proportion of *Vcam1*^+^ sinusoidal endothelial cells (ECs) in young and old skulls and femurs.

b, Representative coronal section images of postnatal day 0, 40 weeks and 80 weeks old mice showing Endomucin (red) expression pattern. Scale bar: 300 μm. c, Representative 3D images (parietal region) from murine young (YS) and old skulls (OS) immunolabelled with nuclear stain DAPI (magenta) and immunostained for FSP1 (green). Bar graph showing quantification of FSP1^+^ bone marrow area (*n* = 5 animals per age group). Scale bar: 20 μm.

d, Representative images (left) of skull marrow (parietal region) from murine YS and OS labelled with nuclear stain DAPI (red) and immunostained for γ-H2AX (green). Bar graphs showing quantification of γ-H2AX^+^ bone marrow area (*n* = 5 animals per age group). Scale bars: 5 μm.

e, Representative images of skull bone marrow (parietal region) from murine YS and OS labelled with nuclear stain DAPI (magenta) and immunostained for P16 (green). Bar graph showing quantification of P16^+^ area (*n* = 5 animals per age group). Scale bar: 20 μm.

Data are mean ± s.d. *P* values by two-tailed unpaired Student’s t-test.

**Extended Data Fig. 2: Age-dependent changes in skull bone and bone marrow**

a, Representative images of skull bone marrow (parietal region) from murine young and old skulls labelled with nuclear stain DAPI (blue) and immunostained for CD3 (green) and CD8 (red). Bar graphs showing quantification of CD3^+^ and CD8^+^ areas (*n* = 5 animals per age group). Scale bars: 20 μm.

b, Representative images of skull marrow (parietal region) from murine young and old skulls labelled with nuclear stain DAPI (blue) and immunostained for perilipin (green). Representative images (right) of skull marrow (parietal region) from murine and old skulls labelled with nuclear stain DAPI (magenta) and BODIPY (green). Bar chart showing quantification of Perilipin^+^ area (*n* = 5 animals per age group). Scale bars: 100 μm (left); 10 μm (right).

c, Representative images of skull marrow (parietal region) from murine young and old skulls labelled with nuclear stain DAPI (blue) and immunostained for VEGFA (red). Representative images of skull marrow (parietal region) from murine young and old skulls labelled with nuclear stain DAPI (blue) and immunostained for OSX (green) and Podocalyxin (red). Scale bars: 20 μm (left); 40 μm (right). Bar charts showing quantification for VEGFA^+^ and OSX^+^ areas (*n* = 5 animals per age group).

d, Representative images of skull marrow (parietal region) from murine young and old skulls labelled with nuclear stain DAPI (blue) and immunostained for Podocalyxin (red) and Ki67 (green). Scale bars: 100 μm (lower magnification); 40 μm (lower magnification). Bar chart showing quantification for Ki67^+^ areas (*n* = 5 animals per age group).

Data are mean ± s.d. *P* values by two-tailed unpaired Student’s t-test.

**Extended Data Fig. 3: Comparison of Endomucin immunofluorescence**

Biologically incorrect nuclear Endomucin immunofluorescence (DOI: 10.1038/ s41586-024-08163-9). Cell surface localised Endomucin immunofluorescence (DOI: 10.1016/j.celrep.2017.01.042).

**Extended Data Fig. 4: 3D imaging of vascular network in interparietal region of murine skull bones**

a, Experimental scheme for thick-sectioning of interparietal region of murine young (8 weeks) and ageing skulls (54 weeks), followed by 3D imaging and vascular analysis.

b, Representative maximum-intensity projections of consecutive sections from the interparietal region of murine young and ageing skulls labelled with nuclear stain DAPI (grey) and immunostained for Endomucin (magenta) and CD31 (green), showing Endomucin+CD31 cells in yellow. BM: bone marrow; Scale bars: 500 μm (overview images); 50 μm (all other images).

**Extended Data Fig. 5: 3D imaging of whole-skulls from mice**

a, Young skull prior to and after tissue clearing. 3D images of the whole skull from *Cdh5 Cre ERT2 X Td-tomato* mice. Scale bars: 500 µm (whole skull); 50 µm (right). b, Whole-mount preparation of murine skull showing nuclear staining with DAPI (blue). Scale bars: 1000 μm (overview images); 300 μm (high-magnification images).

**Extended Data Fig. 6: Ageing drives bone marrow adiposity**

a, Representative images of tibia and vertebra from young and old mice labelled with nuclear stain DAPI (grey) and stained for BODIPY (yellow). Scale bars: 300 µm (overview); 100 µm (high-magnification).

b, Representative 3D images of tibia and vertebra from young and old mice labelled with nuclear stain DAPI (grey) and immunostained for Perilipin (yellow). Bar graphs showing quantification for Perilipin^+^ areas (*n* = 5 animals per age group). Scale bars: 300 µm (overview); 100 µm (zoom-in).

Data are mean ± s.d. *P* values by two-tailed unpaired Student’s t-test.

## Legends to Movies

**Movie 1:** Light-sheet imaging showing skull vasculature.

**Movie 2:** Micro-CT showing young versus old long bones; tibia and femur.

